# Estimating the density of small population of leopard *Panthera pardus* using multi-session photographic□sampling data

**DOI:** 10.1101/2020.08.22.262980

**Authors:** Mohammad S. Farhadinia, Pouyan Behnoud, Kaveh Hobeali, Seyed Jalal Mousavi, Fatemeh Hosseini-Zavarei, Navid Gholikhani, Hasan Akbari, Morteza Eslami, Peyman Moghadas, David W. Macdonald

## Abstract

West Asian drylands host a number of threatened large carnivores, including the leopard *(Panthera pardus)* which is limited to spatially scattered landscapes with generally low primary productivity. While conservation efforts have focused on these areas for several decades, reliable population density estimates are missing. Spatially-explicit capture-recapture (SECR) methodology, incorporating animal movement in density estimates, is widely used to monitor populations of large carnivores. We employed multi-session SECR modeling to estimate the density of a small population of leopard (*Panthera pardus*) in a mountainous stretch surrounded by deserts in central Iran. During 6724 camera trap nights, we detected eight and five independent leopards in 2012 and 2016 sessions, respectively. The top performing model demonstrated density estimates of 1.6 (95% CI = 0.9-2.9) and 1.0 (95% CI = 0.6-1.6) independent leopards/100 km^2^ in 2012 and 2016, respectively. Both sex and season had substantial effects on spatial scale (*σ*), with larger movements for males and during winter. Currently available estimates in arid regions represent some of the lowest densities across the leopard global range. These small populations are vulnerable to demographic stochasticity. Monitoring temporal changes in population density and composition can inform conservation priorities.

## Introduction

The monitoring of large carnivores living in mountainous ecosystems represents a formidable challenge for conservationists and managers. For example, carnivores inhabiting mountainous landscapes tend to persist at lower densities than carnivores in lowland habitats (Alexander et al., 2015; Ghoddousi et al., 2010; Kachel et al., 2017) and have large-scale movement and home ranges (Cheraghi et al., 2019; Farhadinia et al., 2018b; Johansson et al., 2016). Spatially-Explicit Capture Recapture (SECR) models are widely used to monitor populations of large carnivores. The SECR models incorporate spatial locations of captures within a unified model to provide reliable estimates of density (Borchers and Efford, 2008; Efford, 2004). They estimate animal density from a set of individual animal detections made at capture locations, for example by means of motion-detector camera traps, nested within a broader network of potential home-range centres (Efford, 2004).

However, when the population size is very small, several challenges arise for SECR density estimates (Gerber et al., 2014). Most importantly, infrequent detections (Mohamed et al., 2019; Rostro-García et al., 2018), the low number of individuals detected (Hearn et al., 2017; Sharma et al., 2014) and variability in detection among individuals (Alexander et al., 2015; Gerber et al., 2014) can compromise SECR models’ precision. The SECR method also requires sampling to be conducted relatively densely to obtain sufficient recaptures of individuals on multiple cameras; this is required for adequate estimates of the spatial scale parameter (*σ*) (Sollmann et al., 2013; Wilton et al., 2014).

Nonetheless, SECR models perform well for species with large home ranges that are present at low densities if some realistic requirements are met (Wilton et al., 2014; Zimmermann and Foresti, 2016). First, the extent of the detector array has to be similar or larger than the extent of individual movement (Efford, 2011; Sollmann et al., 2012). Second, the optimal distance between traps is twice as large as the *σ* (Sollmann et al., 2012; Sun et al., 2014). Third, repeat detections should exceed 20 (Efford, 2011), and finally, each individual should be detected on average of at least 2.5 times (Gerber et al., 2014). Importantly, sharing detection information across sessions over time and places (Gerber et al., 2014; Morehouse and Boyce, 2016) can increase the precision of density estimates. However, when landscape features restrict the spatial arrangement of detectors, notably in mountainous areas, biologists inevitably select small-sized study areas, usually in good quality habitats, which can result in violating some of the above requirements (Suryawanshi et al., 2019).

Asian mountains harbor large felids with large spatial requirements, such as common leopards (*Panthera pardus*) and snow leopard (*P. uncia*) (Farhadinia et al., 2018b; Johansson et al., 2016). Hence, most population estimates, based on SECR photographic data, come from study areas that are small relative to the species home range (Alexander et al., 2015; Farhadinia et al., 2019; Kachel et al., 2017; Suryawanshi et al., 2019), usually with small populations of fewer than 10 individuals (Ghoddousi et al., 2010; Kachel et al., 2017; McCarthy et al., 2008). Their rarity and potential vulnerability to demographic stochasticity underscores the critical and time-sensitive need to develop e□cient analytical methods so that conservation plans have the greatest chance of success.

In this study, we employed multi-session SECR modeling to understand population density and composition of Persian leopards (*P. p. saxicolor*) in a desert mountain in central Iran. We expected that leopards show inter-seasonal variation in density parameters, with larger movement in winter than summer. Our study provides the density of leopards from the driest area in which leopards have ever been studied globally.

## Materials and Methods

### Study area

The Bafq Protected Area (hereafter Bafq) was designated a protected area in 1996, near the city of Bafq in central Iran. With an area of 885 km^2^, Bafq is an arid mountainous region typified by sparse plains and rolling hills (Figure 1). The altitude range varies between 1060 and 2860 m a.s.l and mean annual precipitation is 70 (Sohrabinia and Hosseini-Zaverei, 2010). Springs, wells, air pumps and small dams constructed at high altitude are the main water supplies of the region.

**Fig 1.**
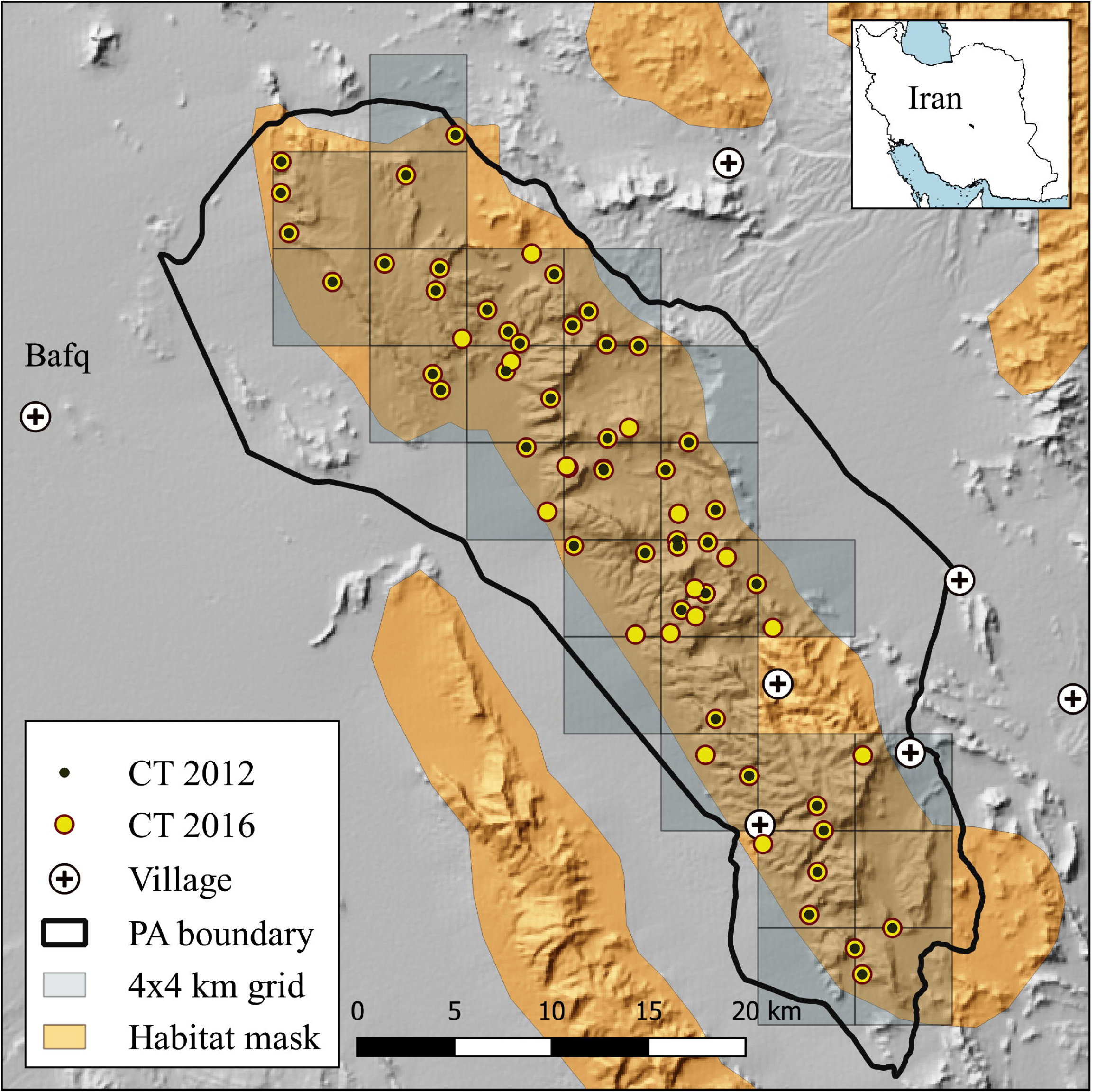
Spatial configuration of study areas and locations of camera trap stations across the two sessions in Bafq Protected Area, central Iran. The map inset shows locations of the study area in Iran.

### Sampling design

We deployed camera trap stations on park-wide 4×4 km grids, ensuring even coverage of the whole area (Figure 1) for 88 and 91 days in winter 2011-2012 and summer-autumn 2016 within Bafq, respectively (Table 1). Short sampling periods (≤ 3 months) are necessary to avoid violating the assumption of demographic closure (Zimmermann and Foresti, 2016). Open desert areas and villages were masked out from the sampling grids. Camera stations were placed at a mean spacing of 1620 and 1409 m in the two consecutive sessions in order to simultaneously achieve the twin objectives of maximizing the number of individuals caught and adequately recapturing individuals at different camera traps, as required in SCR designs. A total of 22 and 26 grids were sampled during the two sessions, respectively (Figure 1).

**Fig 1.**
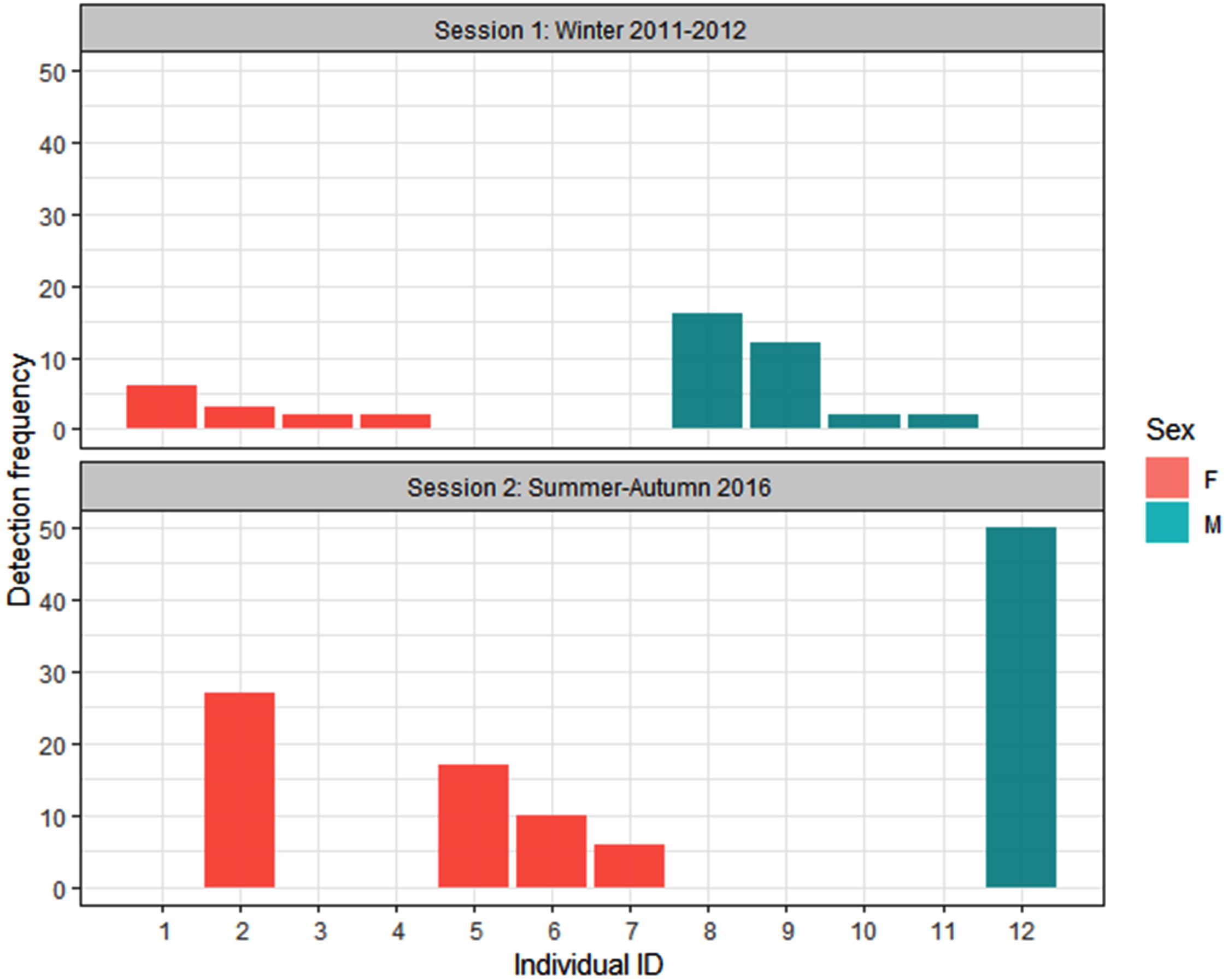
Comparison of detection frequency for all demographic classes between winter 2011-2012 and summer-autumn 2016 sessions in Bafq Protected Area, central Iran. Each code on the x-axis refers to a single individual leopard. Only one individual (F2) was detected in both sessions.

**Table 1.** Details of sampling design for multi-session spatially-explicit capture-recapture framework in Bafq Protected Area (2011-2012 and 2016).

| Session | # stations (# leopard<br>positive stations) | Sampling period (days) | Effort<br>(trap nights) |
| --- | --- | --- | --- |
| Winter 2011-2012 | 47 (19) | 24.12.11 to 13.3.2012 (88) | 2823 |
| Summer-Autumn 2016 | 66 (34) | 27.8 to 26.11.2016 (91) | 3901 |

We deployed Panthera® IV and V (New York, NY 10018, USA) and Cuddeback Capture Model 1125 (Non Typical, Inc., Park Falls, WI, USA), all working with white Xenon flashes. In winter 2011-2012, camera traps were placed either along trails or dirt roads. In 2016, when the survey was conducted during the driest months of year (August-October), we equipped 38 water resources (springs or artificial waterholes, 58% of stations) with a camera trap. We also placed cameras along trails (n=28, 42% of stations), predominantly along valley bottoms or trails.

### Data analysis

We considered each day as a sampling occasion. All consecutive photographs of the same individual taken no more than 0.5 hours apart were defined as a single event. The identity of leopards was determined by the unique rosette patterns on their pelage, independently by two researchers (PB and MSF). Sex was distinguished where possible from sex-specific cues, such as visible genitalia or the presence of dependent individuals.

We estimated density using the maximum likelihood spatially explicit capture– recapture method (Borchers and Efford, 2008) using the package ‘secr’ version 3.2 (Efford, 2019) in the *R* software version 3.3.3. (R Development Core Team, 2013). The estimates of population density produced by SECR are unbiased by edge effects, incomplete detection and heterogeneous capture probabilities and eliminate the need for an ad hoc estimation of the sampling area (Efford, 2004). The secr package also allowed us to evaluate the effect of sex on density parameters.

We combined the two years into a single model framework and considered each year an independent session. For each ‘session’, the secr assumes a new realisation of the underlying population process (Efford, 2019). The multi-session analysis enables the fitting of models with parameter values that apply across sessions, data are then effectively pooled with respect to those parameters. Our two sessions did not overlap temporally and only one individual (F2) was observed in both sessions, ensuring that the independence assumption between the sessions was not violated.

We used a hierarchical model composed of an explicit state-space process model and an observation model (Efford, 2004). The animal population size and their respective central locations (“home-range centers”) constitute the state-space process, assuming a Poisson distribution (Borchers and Efford, 2008; Efford, 2004). The observation model describes the probability of encounter as a function of an individual’s location at the time of sample, and a probability of “count” detector parameter (Efford, 2017). The half-normal detection function contains two parameters. *g*_*0*_ which is the baseline detection rate when the distance between the animal’s activity centers and the camera traps is zero and *σ* which is the spatial scale parameter (with the unit of meters) of the encounter probability model (Borchers and Efford, 2008; Efford, 2004).

We defined the area of integration (i.e., state-space model) by equally spaced points in a regular grid, with a mesh spacing of 1 km^2^. A buffer was plotted around the detector array to incorporate individuals with activity centeres outside of the trapping area, but whose movement range extends into the sampling area (Borchers and Efford, 2008; Efford, 2004). We used a mask with a buffer of 120 km to define the outer limit of the state-space area. This buffer size (3-4 times *σ*) corresponds to an area that animals outside the buffer had a low probability of being detected, and thus were unlikely to influence density estimates (Efford, 2004). Non-habitat areas such as villages, open deserts and sand dunes were masked out from the effective area, based on an ensemble suitability map developed by Ahmadi et al. (Submitted).

For the space model, we fitted 14 *a priori* models with varying effects on *g*_*0*_ and *σ* in a multi-session framework (Table 2). We included a trap-specific behavioral response in the baseline detection model because we expected that the leopard behaviour can change after being detected at a specific trap for the duration of the session (bk). We were also interested in quantifying sex-specific baseline encounter rate (*g*_*0*_) and spatial scale (*σ*), both of which are widely seen in leopard studies (Braczkowski et al., 2016; Goldberg et al., 2015; Rostro-García et al., 2018). We finally fitted session-stratified estimates, meaning that all parameters vary across sessions (i.e., years), by maximising the likelihood. Akaike’s Information Criterion corrected for small samples (AICc) was used to identify the most parsimonious model (i.e., the lowest AICc score) (Burnham and Anderson, 2002). The highest ranked models (all within 2.0 ΔAICc of the top model) were used to estimate leopard density (*D*), detection rate at home range center (*g*_*0*_) and *σ*.

**Table 2.** Model selection results for two sets of 14 fitted multi-session models (totally 28 models) ranked by Akaike’s Information Criterion corrected for small samples (AICc) for 2012 and 2016 Persian leopard density Bafq Protected Area, central Iran. We fitted models using the half-normal detection function. *g*_*0*_ = baseline detection rate at trap location considered as home range center and sigma (*σ*) = spatial scale parameter. Effects on *g*_*0*_ and *σ* included sex (Sex), session (session) and behavioral response (bk).

| Model description | Parameters<sup>a</sup> | AICc | ΔAICc<sup>b</sup> | AICc weight<sup>c</sup> |
| --- | --- | --- | --- | --- |
| $g_0 \sim (bk + session + Sex)$ $\sigma \sim (Sex + session)$ | 7 | 1830.6 | 0.0 | 0.87 |
| $g_0 \sim bk$ $\sigma \sim session$ | 4 | 1835.9 | 5.2 | 0.06 |
| $g_0 \sim (Sex + session)$ $\sigma \sim (Sex + session)$ | 6 | 1837.1 | 6.5 | 0.03 |
| $g_0 \sim (bk + session)$ $\sigma \sim session$ | 5 | 1838.3 | 7.7 | 0.02 |
| $g_0 \sim bk$ $\sigma \sim 1$ | 3 | 1839.7 | 9.1 | 0.01 |
| $g_0 \sim (Sex + bk)$ $\sigma \sim session$ | 5 | 1841.4 | 10.7 | 0 |
| $g_0 \sim 1$ $\sigma \sim (Sex + session)$ | 4 | 1855.7 | 25.0 | 0 |
| $g_0 \sim session$ $\sigma \sim session$ | 4 | 1862.4 | 31.7 | 0 |
| $g_0 \sim Sex$ $\sigma \sim Sex$ | 4 | 1864.5 | 33.9 | 0 |
| $g_0 \sim (Sex + session)$ $\sigma \sim session$ | 5 | 1865.5 | 34.8 | 0 |
| $g_0 \sim 1$ $\sigma \sim Sex$ | 3 | 1875.5 | 44.9 | 0 |
| $g_0 \sim (Sex + session)$ $\sigma \sim 1$ | 4 | 1876.1 | 45.5 | 0 |
| $g_0 \sim Sex$ $\sigma \sim 1$ | 3 | 1884.1 | 53.5 | 0 |
| $g_0 \sim 1$ $\sigma \sim 1$ | 2 | 1886.2 | 55.5 | 0 |

<sup>a</sup> Number of model parameters
<sup>b</sup> Difference in AICc score compared with smallest AICc score.
<sup>c</sup> AICc model weight.

## Results

### Photographic captures

During a sampling period of 2823 trap nights in winter 2011-2012, we obtained a total of 49 independent events from eight independent leopard individuals. Five leopards were photographed on both flanks, two male leopards were photographed only on their right flank and one female leopard accompanied with a single cub was photographed only on its left flank. For analyses, we assumed eight different leopards were photographed, equally from each sex. In summer-autumn 2016, we obtained 141 independent events of leopards during 3901 trap nights (Table 3). Because 58% of camera traps were deployed at water resources in summer 2016, leopards regularly turned around which enabled us to photograph both flanks and to detect five independent individuals. They included only a single adult male and four adult females. We also detected four leopard families with a total of six cubs (ranging 1-2 per family), equally shared between the two sessions.

**Table 3.**
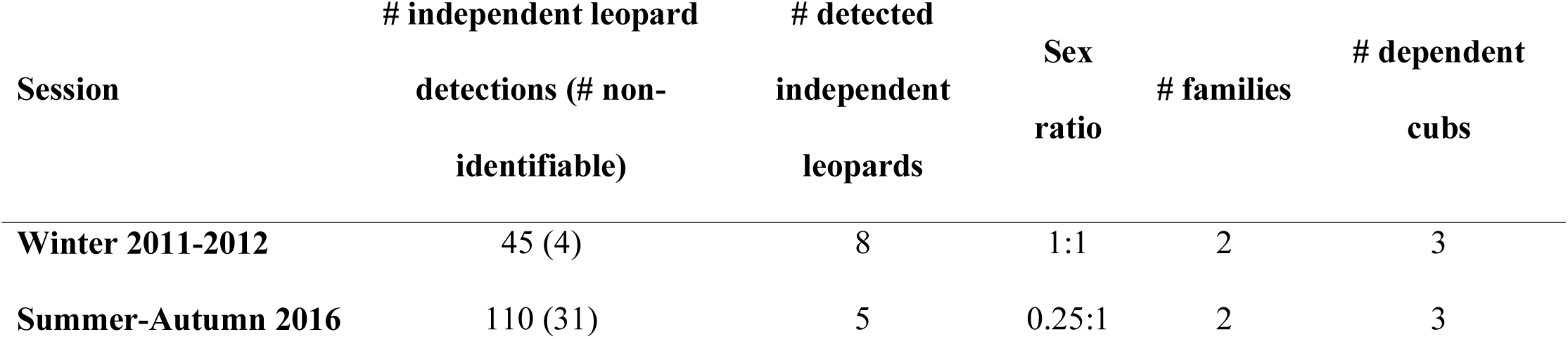
Details of baseline information on leopard based on systematic camera trapping across two sessions of winter 2011-2012 and summer-autumn 2016.

Leopard detections were approximately 3 times more frequent at water-based cameras compared with trail-based stations (105 vs. 36 independent detections; X^2^ = 5.2, df = 1, *P* = 0.02). The four independent females observed during the second session were detected only 1 to 4 times at trail-based camera traps, while they were much more frequently seen at water resources, varying between 5 and 24 detections. During the winter session, same-trap recapture was rare (10% of total detections) whereas it increased to 27% of detections during the summer session. Male leopards showed large scale movements relative to the size of the detector array, with the elongated activity areas aligned with the detector array.

### Density estimation

There was strong support for one model based on the AICc (Table 2), defined as *g*_*0*_ ∼ bk + session + sex *σ* ∼ session + sex. We found sex and session-specific variability in spatial scale (*σ*). Males had larger *σ* than females in each session, yielding an inter-sexual ratio of 1.6 (Table 4), indicating large spatial movements of male leopards throughout Bafq. Likewise, although *σ* estimates were overlapping in their 95% confidence intervals between the two seasons for each sex, the mean estimates of *σ* for the winter session were 1.5 larger than summer-autumn session, regardless of the sex (Table 2). This suggests less mobility of leopards during warm months.

**Table 4.**
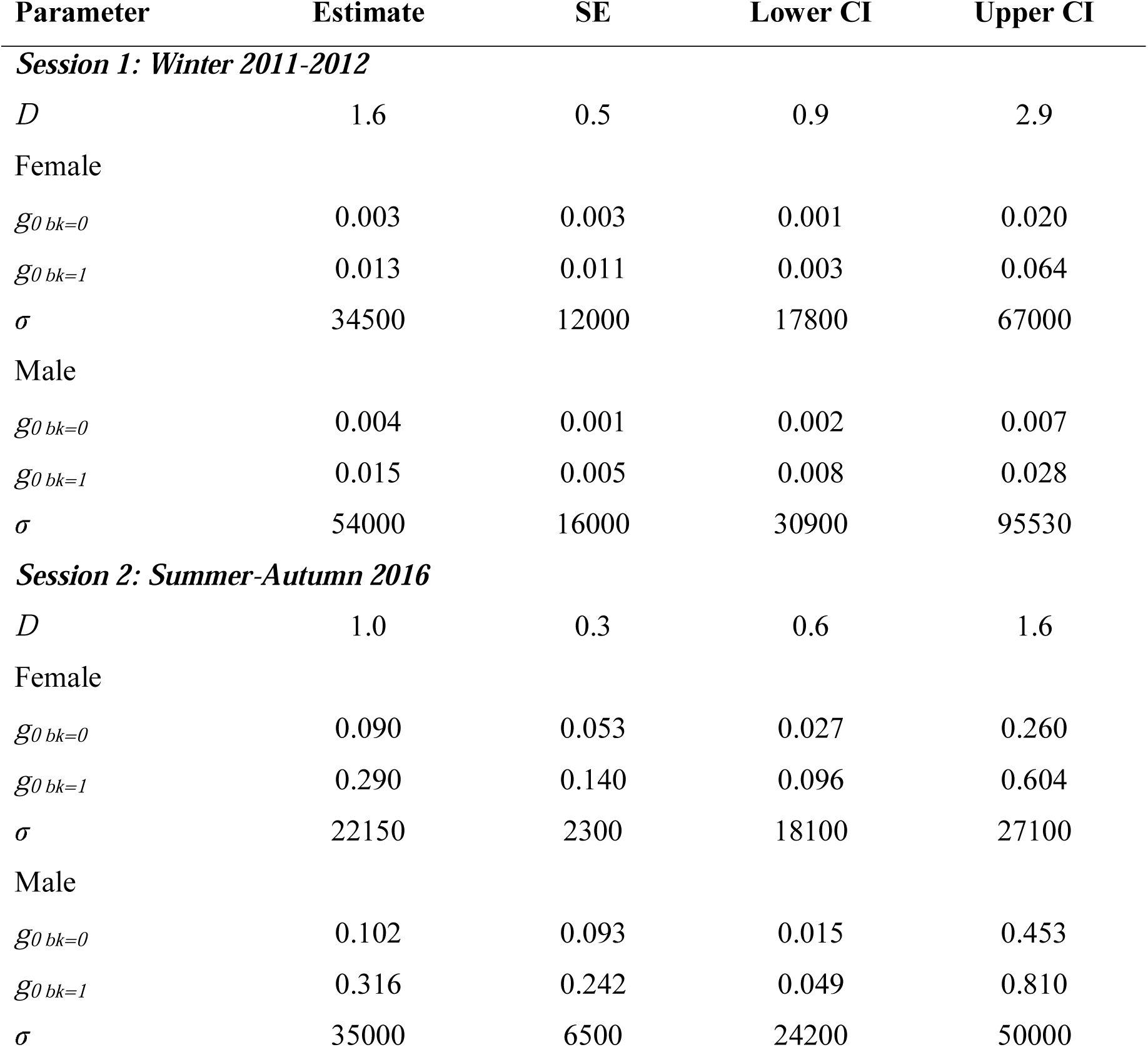
Density estimates of Persian leopards with standard error (SE) and 95% confidence interval (lower and upper) of parameters for spatially-explicit capture recapture models fit to camera trapping data from three study areas in northeastern Iran. Density (*D*) is reported in independent leopards per 100 km^2^. The detection rate at trap location (*g*_*0*_) considered as home range center and *σ* is the scale of an individual’s movement distribution (m). The *g*_*0 bk=0*_ corresponds to the naive state whereas the *g*_*0 bk=1*_ represents learned response.

The top model also supported sex and session-specific baseline detection rate (*g*_*0*_). Thus, the *g*_*0*_ estimates showed substantial inter-sessional difference for each sex, increasing by a magnitude of 21.4-30.0 from the winter to summer-autumn sessions (Table 2). For all sex groups within a session, *g*_*0 bk=1*_ was larger than *g*_*0 bk=0*_ with a ratio of 3.2-4.3 between the means. The bk effect tests the hypothesis that leopard behaviour changes after being detected at a specific site for the duration of the survey (trap response). Therefore, when a leopard is detected in a specific camera trap site, the probability of a subsequent encounter for the entire survey is increased, i.e. the individual becomes ‘trap happy’ at the population level (beta coefficient _g0.bk_=1.4, 95% CI = 0.9-1.8).

The best performing model estimated a density of 1.6 (95% CI = 0.9-2.9) and 1.0 (95% CI = 0.6-1.6) independent leopards/100 km^2^ for the first and second sessions, respectively (Table 4). A total of 10.9 ± SE 3.4 independent leopards were estimated for the winter 2011-2012 while the population size was calculated as 6.9 ± SE 1.8 independent leopards for the 2016 session.

## Discussion

We estimated the density of leopards from the driest area in which leopards have ever been studied globally, with an average annual precipitation of 70 millimetres (Sohrabinia and Hosseini-Zaverei, 2010). Although data on leopard density estimates in arid regions are sparse, they, including that from this study site, indicate some of the lowest densities across the leopard global range, all below 2 adults per 100 km^2^ (Edwards et al., 2016; Ghoddousi et al., 2010; Stein et al., 2011).

Our analysis did not support a “density variation”, i.e. real population differences, between the two sessions as the confidence intervals overlapped substantially. Alternatively, the different density point estimates could be “sampling variation”, i.e. variation in a statistic from sample to sample, which is commonly seen in multi-session SECR studies of leopards (Rogan et al., 2019; Rosenblatt et al., 2016). There are two possible reasons supporting that the density has not changed between the two sessions. First, “top-down” regulation due to human persecution which causes higher anthropogenic mortality (Rosenblatt et al., 2016; Rostro-García et al., 2018; Sharma et al., 2014). However, we found little evidence that the Bafq leopard population suffers direct anthropogenic persecution. As such, only one record of an individual poisoned between the two survey efforts was obtained. Importantly, conflict with communities, which is a major source of concern in many parts of the leopard range in the Middle East (Babrgir et al., 2017; Kabir et al., 2013), seems to be limited; a recent dietary investigation based on faecal analysis found only 7.54 % of biomass consumed composed of domestic animals in Bafq (Rezaei et al., 2016).

Alternatively, prey density as the main ‘bottom-up’ process can shape demography in leopards (Henschel et al., 2011; Ramesh et al., 2017). We obtained annual census data for the period between 2011 and 2016, conducted by Yazd Provincial Office of Department of the Environment every November using 10-15 groups of field crew (Table S1), suggesting that the population size for bezoar goat and urial, two key prey for Persian leopards (Farhadinia et al., 2018a; Rezaei et al., 2016), was not decreasing. Therefore, there is little evidence that the bottom-up process was in charge of demographic changes in Bafq. However, small sample sizes and a lack of more replications require us to view our findings as suggestive rather than conclusive, and to concede that further research is necessary to assess the demographic trends of leopards in central Iran.

Importantly, season had a strong effect on baseline detection rate (*g*_*0*_) and spatial scale (*σ*), both of which influence the density estimate. Nonetheless, the season can have confounding effects with year. Larger *σ* during winter (2011-2012) suggested higher mobility of leopards preceding and during mating season, which peaks in mid-winter (Farhadinia et al., 2009). In contrast, water resources are crucial for leopards in summer, not only to meet their drinking requirements, but to optimize their encounter rate with prey (Farhadinia et al., 2018a). This results in smaller *σ* and larger *g*_*0*_ in warm season. In addition to season, leopard populations may exhibit marked inter-annual variation in *σ* (Rogan et al., 2019; Rosenblatt et al., 2016), possibly due to their dynamic spatial patterns and home range sizes.

The persistence of small leopard populations can be adversely affected by demographic stochasticity through reducing survival and fecundity (Rostro-García et al., 2018; Williams et al., 2017). Importantly, controlling two threats, prey depletion and leopard persecution, both inside Bafq as well as nearby areas to support individuals dispersing from this breeding population are crucial. In addition to single density estimates, temporal changes in population density and composition are helpful to inform data-driven decision-making process to define conservation priorities. Yet importantly, to the best of our knowledge, no study has assessed demographic trends of leopards, and other large carnivores inhabiting remote drylands of the Middle East. Addressing this knowledge gap is essential for leopard management and conservation.

## Acknowledgements

We sincerely thank the Iranian Department of Environment for administrative support and provision of necessary permissions. Financial support was provided by Conservation of Asiatic Cheetah Project, Yazd Provincial Office of Department of the Environment, Dieren Park Amersfoort Wildlife Fund and Rufford Foundation. E. Horstman and A. Morehouse advised on data analyses and P. Johnson, L. Sousa and A. Braczkowski commented on an earlier version of this manuscript. A. Rezaei, A. Bakhtiari and H. Alizadeh assisted in field sampling. We are particularly grateful to the support and dedication of the Bafq’s rangers.

## Data availability

Datasets analysed during the current study are available on Figshare with the generated link as (https://figshare.com/s/98ca1dc84ffc861dbed7).

