## Supplemental Table 1 for "Estimating the density of small population of leopard *Panthera pardus* using multi-session photographic□sampling data"

Table S1 Population size of mountainous ungulates in Bafq Protected Area, based on annual census carried out every November using 10-15 groups of field crew (data from Yazd Provincial Office of Department of the Environment). No census was conducted in 2012 and 2016 due to unpredictable weather conditions.

| **Year** | **Urial** | **Bezoar goat** |
| --- | --- | --- |
| 2010 | 316 | 398 |
| 2011 | 274 | 765 |
| 2012 | NA | NA |
| 2013 | 229 | 688 |
| 2014 | 207 | 781 |
| 2015 | 231 | 957 |
| 2016 | NA | NA |
| 2017 | 631 | 922 |
